# *In vivo* imaging of calcium dynamics in zebrafish hepatocytes

**DOI:** 10.1101/2022.01.11.475536

**Authors:** Macarena Pozo-Morales, Inés Garteizgogeascoa, Camille Perazzolo, Sumeet Pal Singh

## Abstract

Hepatocytes were the first cell-type for which oscillations of cytoplasmic calcium levels in response to hormones were described. Since then, investigation of calcium dynamics in liver explants and culture has greatly increased our understanding of calcium signaling. A bottleneck, however, exists in observing calcium dynamics in a non-invasive manner due to the optical inaccessibility of the mammalian liver. Here we take advantage of the transparency of the zebrafish larvae to develop a setup that allows *in vivo* imaging of calcium flux in zebrafish hepatocytes at cellular resolution. Using this, we provide quantitative assessment of intracellular calcium dynamics during multiple contexts, including growth, feeding, ethanol-induced stress and cell ablation. Specifically, we show that synchronized calcium oscillations are present *in vivo*, which are lost upon starvation. Feeding recommences calcium waves in the liver, but in a spatially restricted manner. Further, ethanol treatment as well as cell ablation induces calcium flux, but with different dynamics. The former causes asynchronous calcium oscillations, while the latter leads to a single calcium spike. Overall, we demonstrate the presence of oscillations, waves and spikes *in vivo*. Thus, our study introduces a platform for observing diverse calcium dynamics while maintaining the native environment of the liver, which will help investigations into the dissection of molecular mechanisms supporting the intra- and intercellular calcium signaling in the liver.

## INTRODUCTION

Agonist-induced oscillations in cytoplasmic calcium levels were first described for hepatocytes in 1986 (Woods et al., 1986). Since then, the exploration of calcium dynamics and signaling in the liver has contributed considerably to our knowledge of calcium signaling. These studies have positioned intracellular calcium as a key part of the information processing network, wherein environmental inputs are connected to cellular response by calcium ions acting as a key secondary messenger. As an important part of signal integration and pathway crosstalk, intracellular calcium regulates a multitude of physiological processes in hepatocytes, including cell-cycle, function such as bile secretion and glucose metabolism, response to cell death, and regeneration (Amaya and Nathanson, 2013; Gaspers et al., 2012; Nathanson and Schlosser, 1996; Oliva-Vilarnau et al., 2018).

Our understanding of the role of calcium in the liver has been greatly aided by studies in cultured hepatocytes, liver slices and explants. Though the *ex vivo* setup allows precise control of environmental factors, it, however, cannot completely mimic the native environment. Recently, efforts have been made to image liver calcium dynamics *in vivo*. For instance, Lin and colleagues developed bioluminescence based non-invasive imaging of intracellular calcium levels for mouse liver (Oh et al., 2019). Though intracellular calcium changes for an entire lobe could be successfully recorded, the lack of optical accessibility to the mouse liver made it impossible to achieve a higher resolution. Averaging of calcium signals across the entire lobe misses key information related to the temporal and spatial pattern of inter-cellular calcium flux. As the dynamic pattern of intracellular calcium level encodes information related to the biological role of the secondary messenger (Berridge, 1997; Brodskiy et al., 2019; Noren et al., 2016), a window into calcium dynamics *in vivo* at cellular resolution would help to improve our understanding of the role of calcium in liver physiology.

Here we accomplish non-invasive imaging of liver calcium dynamics at cellular resolution using the zebrafish model system. Zebrafish liver is similar to mammalian liver in terms of cellular composition, transcriptional profile and functionality (Morrison et al., 2021). In addition, zebrafish provide unique opportunities in imaging due to external development and transparency of the larval zebrafish. Here, we take advantage of these features to develop an ideal tool for *in vivo* imaging of calcium dynamics. We achieve this by generating a new transgenic model expressing a sensitive genetically encoded calcium sensor specifically in the zebrafish hepatocytes. With this, we provide quantitative assessment of calcium dynamics during multiple physiological contexts, setting up a platform for identifying and exploring the mechanistic role of calcium signaling in the liver.

## RESULTS AND DISCUSSION

### A setup for cellular resolution imaging of cytoplasmic calcium in zebrafish hepatocytes

To visualize the intracellular calcium in zebrafish hepatocytes, we generated a new transgenic line expressing GCaMP6s, a highly-sensitive genetically encoded calcium sensor (Chen et al., 2013), under the control of *fabp10a* regulatory sequences (Her et al., 2003). Using heterozygote *Tg(fabp10a:GCaMP6s)*, we developed a protocol for confocal imaging of calcium dynamics in zebrafish liver. Here, the zebrafish larvae are immobilized in low-melt agarose and placed with the left-side facing the confocal lens. This allowed us to capture GCaMP6s fluorescence in the hepatocytes within the left lobe of zebrafish larvae (for simplicity, ‘zebrafish liver’ refers to the left lobe of the zebrafish larvae in this manuscript).

During the initial stages of experimental setup, we encountered difficulties in imaging calcium dynamics. Specifically, the liver lacked any calcium dynamics. Upon literature survey of the neuronal calcium-imaging field, particularly studies related to zebrafish epilepsy, we realized that tricaine methanesulfonate (MS-222), the commonly utilized anesthesia for zebrafish, suppresses intracellular calcium oscillations (Turrini et al., 2017). MS-222 blocks sodium channels and reduces action potentials in neurons (Carter et al., 2011). Further, MS-222 has been demonstrated to induce oxidative stress response in fish liver (Velisek et al., 2011). As an alternative to MS-222, tubocurarine, anti-nicotinic neuromuscular blocker, has been previously utilized to immobilize zebrafish larvae for imaging (Favre-Bulle et al., 2018; Winter et al., 2017; Zdebik et al., 2013). When comparing calcium dynamics in the liver between MS-222 or tubocurarine treated zebrafish, we observed robust suppression of calcium oscillations in MS-222 treated animals (Fig. S1, Movie 1). Thus, we utilized the neuromuscular blocker tubocurarine to immobilize zebrafish larvae for all of our experiments.

Immobilized zebrafish were imaged using 3D time-lapse confocal microscopy for a period of 15 minutes. At 5 day post-fertilization (dpf), our setup could robustly capture increase in cytoplasmic GCaMP6s fluorescence, indicating increases in intracellular calcium levels (Fig. 1A, Movie 2). We observed calcium oscillations in individual hepatocytes (Fig. 1B). Moreover, the liver displayed waves of synchronized calcium flux (Fig. 1C). Such calcium waves have been previously described for the mammalian liver (Verma et al., 2018).

**Figure 1:**
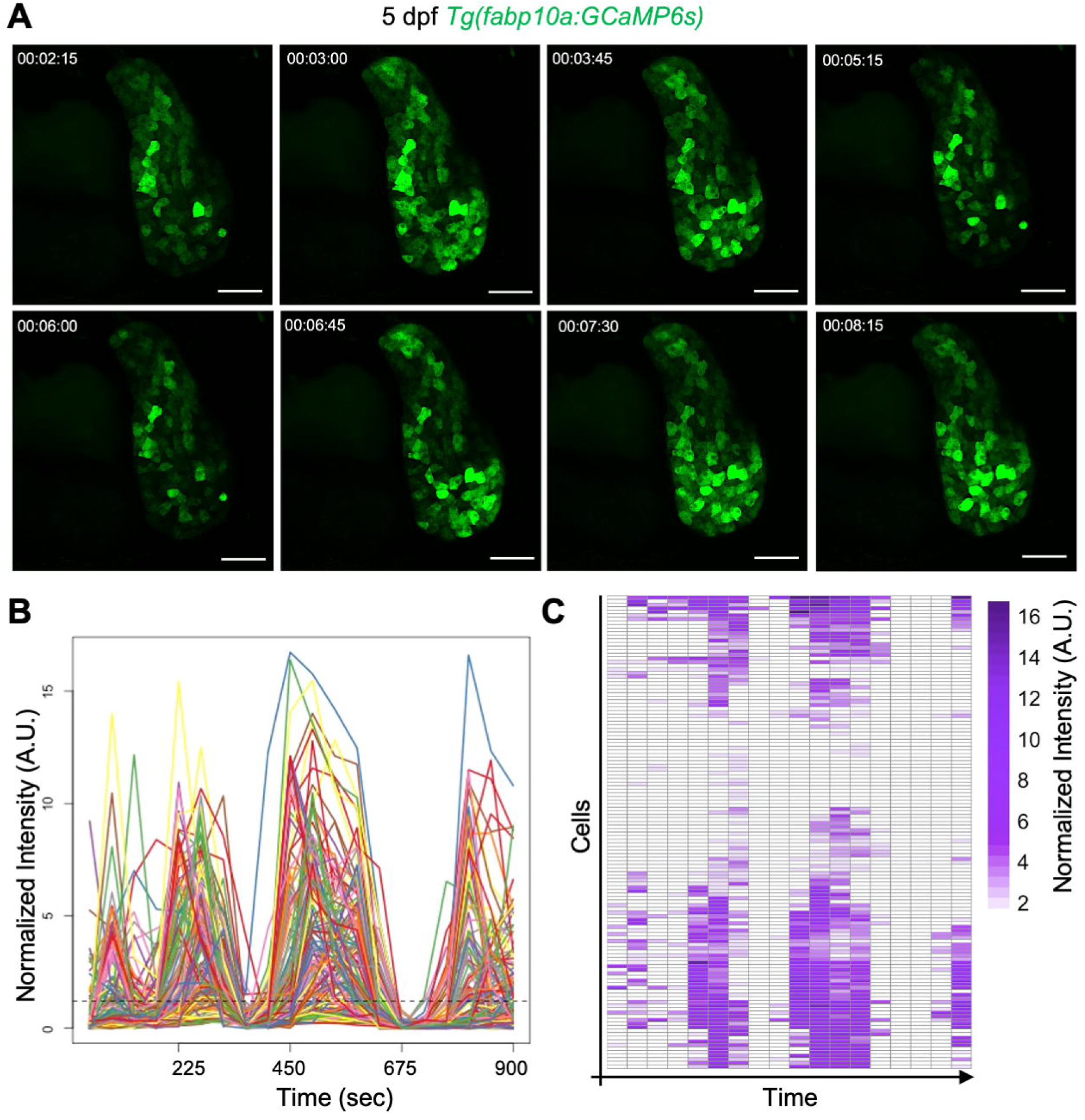
*In vivo* imaging of calcium dynamics in zebrafish hepatocytes at cellular resolution. **(A-C)** Synchronized and repetitive calcium oscillations in the liver. Confocal images **(A)**, line trace **(B)** and heatmap **(C)** are shown for a representative liver of a 5 dpf *Tg(fabp10a:GCaMP6s)* animal. **(A)** Snapshots of liver from time-lapse imaging. The hepatocytes show periodic changes in intracellular calcium levels spaced by approximately four minutes. Scale-bar: 50 µm. **(B)** Line trace of normalized GCaMP6s fluorescence intensity plots of the liver. Each trace corresponds to a single hepatocyte. **(C)** Heatmap visualization of normalized GCaMP6s fluorescence intensity. Each row in the heatmap (Y-axis) represents a single hepatocyte, while every column (X-axis) corresponds to a single frame of time-lapse imaging.

Overall, we developed a system for cellular-resolution visualization of calcium flux in the hepatocytes of alive zebrafish, providing us with a unique opportunity to investigate the phenomenon in the native environment.

### Calcium oscillations in zebrafish liver are linked to nutrient availability

Using our setup, we performed a time-course experiment to investigate the liver’s calcium dynamics during growth. Formation of a functional liver in zebrafish is complete between 4 - 5 dpf (Wang et al., 2017), following which the liver grows in size primarily by proliferation of differentiated cells. We characterized calcium dynamics of the liver following functional maturation, from 5 - 7 dpf (Movie 3). The time-course analysis demonstrated the total number of oscillations at 5 dpf are higher than at 6 and 7 dpf (Fig. 2A).

**Figure 2:**
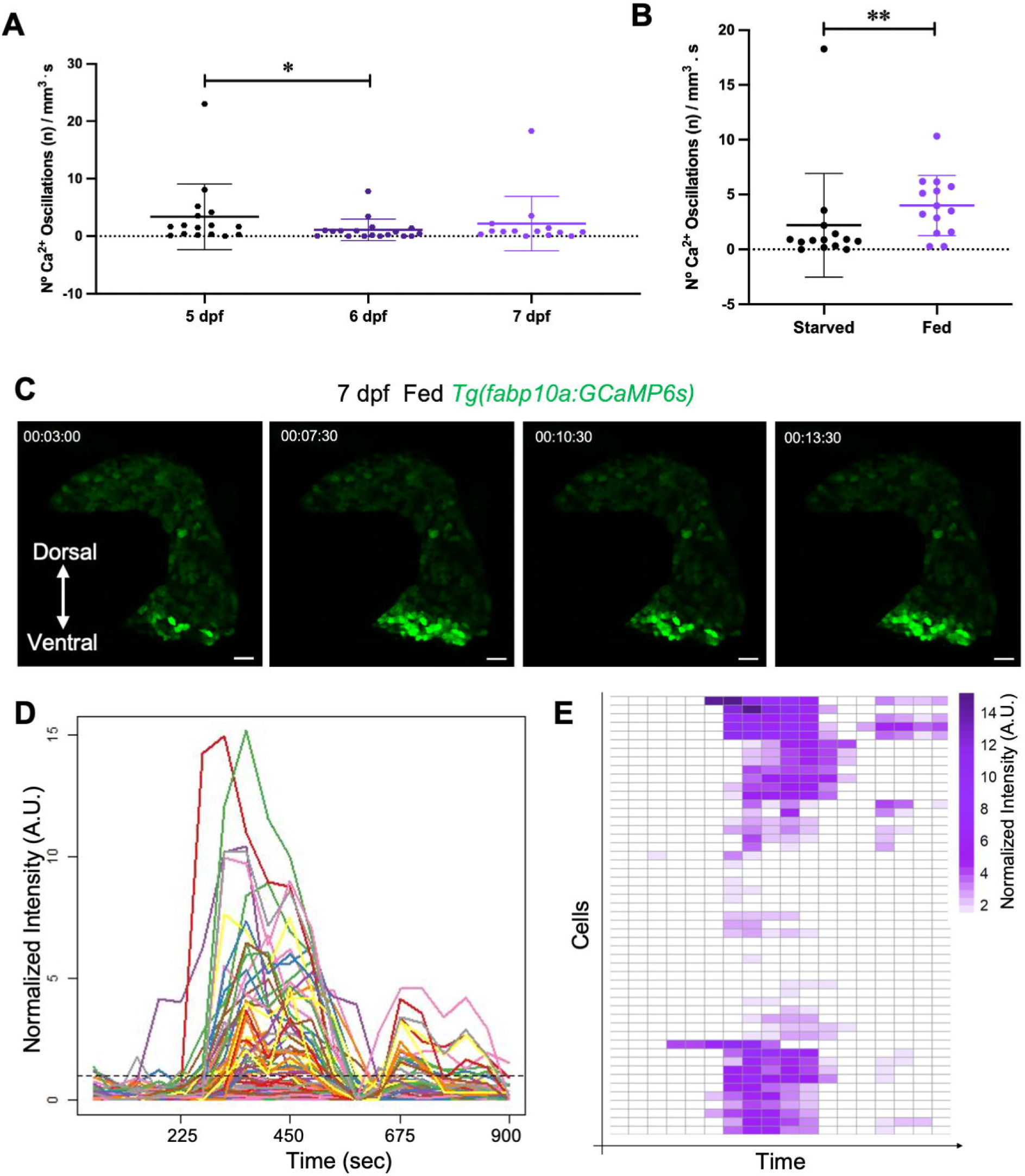
Calcium dynamics during growth and in response to feeding. **(A)** Normalized calcium oscillations at 5, 6 and 7 dpf. Each point represents an individual animal. Mean ± SD is added to the dot-plot. * p-value < 0.05, Mann-Whitney U Test. **(B - D)** Feeding induces spatially-localized synchronized calcium oscillations. **(B)** Dotplot with Mean ± SD showing comparison of calcium oscillations between the ventral half of fed and fasted animals. Fed animals were provided with rotifers from 5 - 7 dpf. ** p-value < 0.01, Mann-Whitney U Test. **(C)** Snapshots from time-lapse imaging of liver from fed *Tg(fabp10a:GCaMP6s)* animal. The ventral tip of the liver demonstrates calcium dynamics. Scale-bar: 50 µm. Line trace **(D)** and heatmap **(E)** representation of normalized fluorescence intensity from 7 dpf fed *Tg(fabp10a:GCaMP6s)* animal.

We hypothesized that the suppression of calcium oscillations in older larvae could be attributed to lack of nutrients. Zebrafish larvae depend on yolk for nourishment, which is depleted between 4 - 5 dpf (Gut et al., 2013). Subsequently, zebrafish larvae display a feeding to fasting transition and enter a starvation state. To test if calcium oscillations in the liver depend on nutrient availability, we fed larvae starting at 5 dpf, which is the standard feeding schedule for zebrafish rearing (Lawrence et al., 2016). At 7dpf, the livers of fed animals displayed synchronized calcium oscillations (Fig. 2D, E). Interestingly, the majority of oscillating cells were localized in the ventral tip of the liver (Fig. 2C, Movie 4). Such spatial-localization was observed in 10 / 14 animals, while only 2/14 animals displayed oscillations in the ventral and dorsal tip of the liver. Further, the ventral region of liver from fed zebrafish displayed significantly more oscillations than livers of fasted zebrafish (p = 0.008) (Fig. 2B).

Our time-course analysis suggests a close relationship between nutrients and calcium signaling in the liver. The two could potentially be linked via insulin / insulin-like growth factor pathway (Rodrigues et al., 2008).

### Ethanol exposure and cellular injury modulate intracellular calcium levels, albeit with different dynamics

Liver damage represents a wide-spectrum of disorders that can progressively lead to liver failure. Understanding the signals initiated during early stages of liver damage could improve disease outcome. To investigate calcium dynamics upon liver damage, we utilized two models: stress model using acute exposure to ethanol (EtOH), and injury model using cell ablation.

In the first model of liver damage, acute stress was induced by exposing zebrafish larvae to 2 % EtOH from 5 dpf for a period of 18 - 22 hours (Fig. 3A) (the four hour time difference in treatment represents the period required for imaging animals present in one replicate). Quantification of calcium oscillations between controls and treated animals demonstrated that acute exposure to EtOH induces calcium oscillations (p = 0.006) (Fig. 3B, Movie 5), suggesting induction of calcium signaling in response to EtOH-induced stress. Further, within the liver, hepatocytes displayed asynchronized calcium oscillations (Fig. 3C, Movie 5).

**Figure 3:**
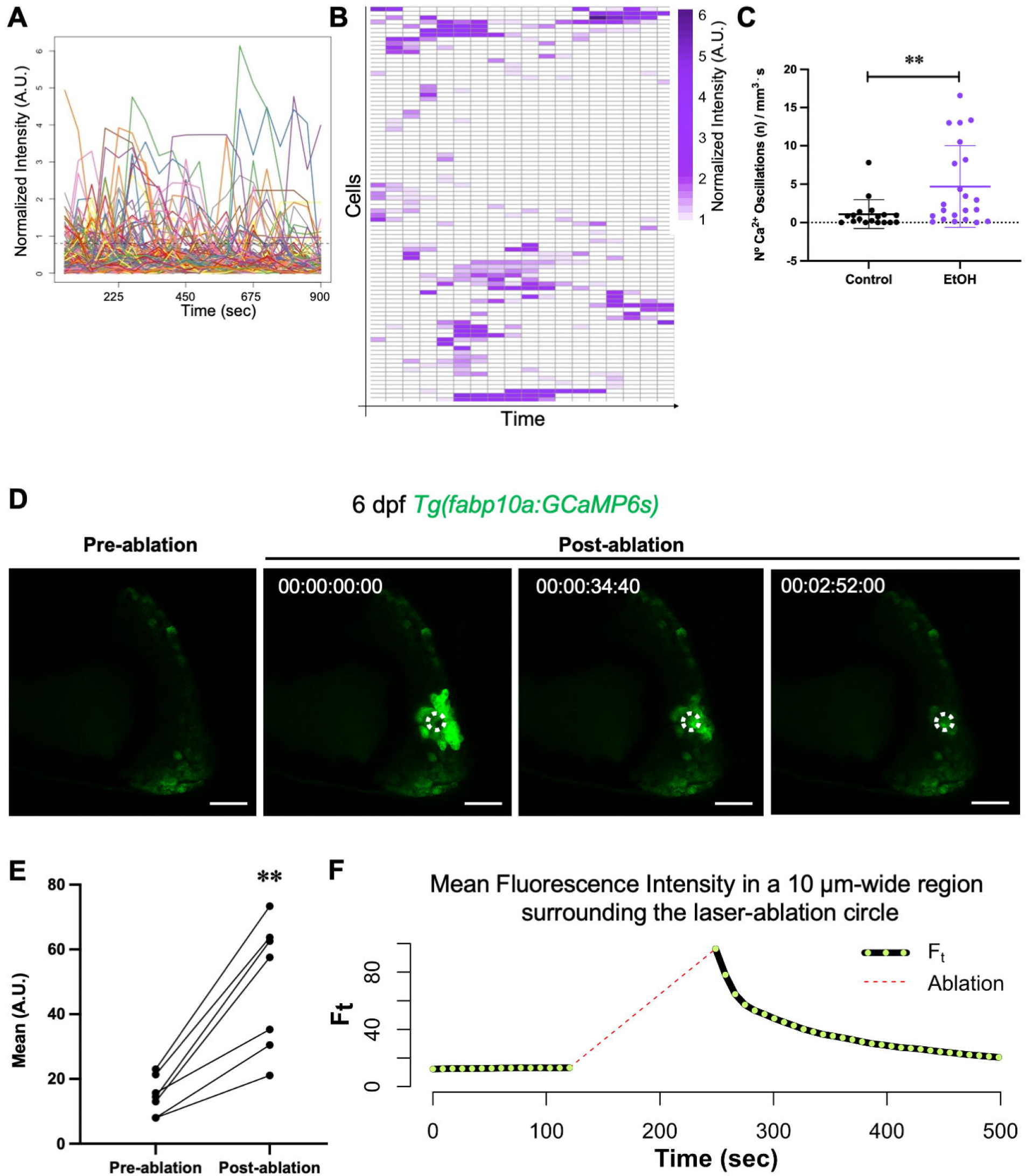
Calcium dynamics in response to ethanol stress and two-photon laser ablation-induced injury. **(A - C)** Ethanol induces asynchronous calcium oscillations. 6 dpf *Tg(fabp10a:GCaMP6s)* animals were imaged after treatment with control or 2 % ethanol for 18 - 22 hours. Line Trace **(A)** and **(B)** heatmap representation of normalized intensity upon ethanol treatment for a representative animal. **(C)** Dotplot with Mean ± SD showing comparison of calcium oscillations between control and ethanol treated animals. ** p-value < 0.01, Mann-Whitney U Test. **(D - F)** Cell ablation induces a calcium spike. Livers from 6 dpf *Tg(fabp10a:GCaMP6s)* were subjected to two-photon laser ablation **(D)** Snapshots from time-lapse imaging of the liver before and after laser ablation. The ablated region is outlined with a dotted white line. **(E)** Dotplot representing the mean fluorescence intensity in the cells surrounding the ablation region. The intensity recorded from individual animals pre- and post-ablation are connected. ** p-value < 0.01, paired t-test. **(F)** Line Trace displaying the mean fluorescence intensity in the cells surrounding the ablation region. The trace corresponds to the representative example. Green dots representing intensity measurements are connected by solid line. Dotted red line corresponds to the time during which two-photon laser ablation was performed. During this time, no readings were obtained.

In the second model of damage, cells in the zebrafish liver were laser-ablated. For this, 6 dpf *Tg(fabp10a:GCaMP6s)* animals were immobilized and the calcium dynamics in the liver were imaged. Following this, a circular region of width 20 µm was selected within the center of the liver and damaged by the two-photon laser ablation. Subsequently, the calcium dynamics were imaged. With this, we observed robust increase in intracellular calcium levels in the cells surrounding the damaged area (Fig. 3D, E, Movie 6). However, cytoplasmic calcium in the neighboring cells did not oscillate, but displayed a strong increase that waned with time, indicative of a single calcium spike (Fig. 3F, Movie 6).

Together, the two models of liver damage implicate calcium signaling in the damage-response process. However, the calcium dynamics differ in the two models, which could be related to the dissimilarity in the damage between the two contexts. The model of acute EtOH stress has been shown to robustly induce unfolded protein response (UPR), ER-stress and accumulation of lipids (steatosis or fatty liver) in zebrafish liver (Lai et al., 2018; Tsedensodnom et al., 2013). Here, calcium signaling could interact with the metabolic rewiring of the hepatocytes, which would be of great interest to dissect. In the model of cell ablation, the damaged and dying cells could release ATP in the extracellular space, where it can activate purinergic receptors in the neighboring cells (Dubyak and el-Moatassim, 1993). The non-oscillatory, but transient, nature of calcium activation in response to ablation might represent the release and clearance of ATP from the extracellular space. Of note, the dynamics of clearance of ATP, and its metabolized products, might differ between *in vivo* and *ex vivo* conditions. This necessitates investigating the damage response in native environment, which is made possible by our setup.

### A platform to investigate hepatocyte calcium dynamics in native environment

In conclusion, we have demonstrated the potential of the zebrafish model system to investigate temporal changes in hepatocyte calcium levels in alive animals. We present the utility of the system under multiple contexts: growth, feeding, stress and injury. Our results raise multiple questions: why are calcium oscillations spatially restricted upon feeding? ; What is the similarity and differences in calcium signaling during growth and EtOH stress? ; What is the role of a transient non-oscillatory calcium signal in the proximity of ablated hepatocytes? These questions, which can be addressed in future, were raised by documenting liver calcium dynamics *in situ*. Thus, our platform offers a valuable tool for documenting the activity of a critical signaling pathway in a wide range of physiological contexts.

## MATERIALS AND METHODS

### Zebrafish strains and husbandry

Wild-type or transgenic zebrafish of the outbred AB strain were used in all experiments. Zebrafish were raised under standard conditions at 28 °C. Animals were chosen at random for all experiments. Zebrafish husbandry and experiments with all transgenic lines will be performed under standard conditions as per the Federation of European Laboratory Animal Science Associations (FELASA) guidelines (Aleström et al., 2020), and in accordance with institutional (Université Libre de Bruxelles (ULB)) and national ethical and animal welfare guidelines and regulation, which were approved by the ethical committee for animal welfare (CEBEA) from the Université Libre de Bruxelles (protocols 578N-579N).

### Generation of *Tg(fabp10a:GCaMP6s; cryaa:mCherry)* zebrafish line

To generate the fabp10a:GCaMP6s; cryaa:mCherry construct, a fabp10a (fatty acid binding protein 10a, liver basic) promoter-containing vector was digested with NheI / NotI. GCaMP6s construct was obtained from the plasmid ins:GCaMP6s; cryaa:mCherry (Singh et al., 2017) using SpeI / NotI. The two were ligated to yield the final construct. The final construct contains fabp10a promoter driving GCaMP6s and a directionally inverted eye marker cassette using a crystallin promoter (cryaa:mCherry) to select transgenic carriers. The entire construct was flanked with I-SceI meganuclease sites to facilitate transgenesis. Transgenic animals were identified by fluorescent red eyes at 3 dpf for experiments. The transgenic line is abbreviated as *Tg(fabp10a:GCaMP6s)* in the manuscript.

### Ethanol Treatment

Transgenic *Tg(fabp10a:GCaMP6s)* animals were treated with 2 % EtOH dissolved in the embryo (E2) medium. 25 larvae were treated in 25 ml of medium (density 1 animal / ml). During the treatment, the petri dish was sealed with parafilm to avoid ethanol evaporation. Treatment was started at 5.25 dpf (afternoon of 5 dpf). On the following day, after a period of 18 hours, imaging was initiated. All replicates were imaged within four hours. As control, non-treated animals were used.

### Confocal Imaging and Analysis

Animals were anesthetized in 0.02% of tricaine methanesulfonate (MS-222) (Sigma, E10521) (Fig. S1) or immobilized in 4 mM of (+)-Tubocurarine chloride pentahydrate (abbreviated as tubocurarine) (Sigma, 93750) and mounted in 1 % Low-Melt Agarose (Lonza, 50080) and imaged on a glass bottom FluoroDish™ (WPI FD3510-100) using a Zeiss LSM 780 confocal microscope. The liver was imaged using a 40x/1.1N.A. water correction lens. Imaging frame was set at 1024 × 1024 pixels and the distance between confocal planes was set at 6 μm for Z-stack to cover on average a thickness of 90 μm. A time-lapse of 20 stacks was conducted with an interval of 45 seconds between each stack. The samples were excited with 488 nm for GCaMP6s, and fluorescence was collected in the range of 499 - 579 nm.

Fiji (Schindelin et al., 2012) was utilized for image analysis using maximum-intensity projections of the Z-stack. In the maximum intensity projections, the cytoplasm of individual cells was delimited manually using the ROI Manager in Fiji. Using the ROI, the mean gray value of, representing the fluorescent intensity of GCaMP6s was extracted. The intensity values for all ROIs in a single image were saved as a comma separated file for downstream processing.

Analysis of intensity values was performed in R (R Core Team, 2020) version 4.0.3. First the intensity values were normalized using the formula:

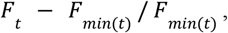

where *F*_*t*_ represents the intensity at a given time, and *F*_*min*(*t*)_ represents the local minima. Local minima is defined as the minimum intensity value between the current frame and two adjacent frames on either side of the time-lapse. The local minima was calculated using the runmin() function from the caTools package (Tuszynski, 2020). Using the normalized intensity values, peaks for the time-lapse were identified using the detect_localmaxima() function from the scorepeak package (Ochi, 2019). The function was utilized from the third to eighteenth frame as local minima could be calculated for those frames (the first two and last two frames do not have two adjacent frames on one side). The detect_localmaxima() function identifies a peak as a signal that shows a local maxima, meaning that the value of the signal rises and then falls. The peaks represent calcium oscillations. Oscillations for each cell were added to obtain a total number of oscillations per image.

For comparison, the normalized number of oscillations was calculated to correct for the size of the liver. For this, volume was measured using Fiji. Here, for each frame in a single Z-stack, the outline of the liver was manually drawn. Then the area of the liver in each frame was extracted. Volume was calculated as (sum of the area) * (thickness of a single Z-stack = 6 μm). This provided volume in μm^3^, which was converted to mm^3^ for normalization. Oscillations were normalized using the formula:

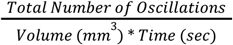

For representation, Fiji was used to add scale bars and PowerPoint was used for adding arrows and labels.

### Statistical Analysis

Statistical analysis was performed using GraphPad Prism version 9.3.1 for Mac OS X, GraphPad Software, San Diego, California USA, www.graphpad.com. The test utilized for comparison is mentioned in figure legend of respective graphs. For all analysis, data was tested for normal distribution. For non-normal distributed data (or if the test for normal distribution could not be performed due to small sample sizes), a nonparametric test was used. No data was excluded from analysis. Blinding was not performed during analysis.

### Two-photon Laser Ablation, Imaging and Analysis

Larvae at 6 dpf from *Tg(fabp10a:GCaMP6s; cryaa:RFP)* were subjected to two-photon laser-induced injury using Zeiss LSM 780 equipped with an objective (40x/1.1 N.A.). For this, the embryos were anaesthetized with tubocurarine and mounted in 1 % low-melting agarose. Single-photon confocal imaging was conducted prior to laser-ablation. For this, a region of four confocal planes near the center of the liver was imaged using a 40x/1.1N.A. Water lens. Imaging frame was set at 1024 × 1024 pixels and the distance between confocal planes was set at 6 μm. Each Z-stack consisting of four frames took 8.6 seconds to image. A time-lapse of 16 stacks was conducted with an interval of 1 ms to allow continuous signal acquisition. Following this, a designated circular area of 20 µm diameter in the center of the Z-stack (single-plane) was exposed to pulsed two-photon laser using Chameleon Vision II (Coherent) at the output power of 2.5 W (λ = 980 nm) for approximately 2 min. Following laser-ablation, single-photon confocal imaging was resumed. Following laser-ablation, 30 stacks were imaged.

For image analysis, the time-lapse was opened in Fiji and a maximum intensity projection image was generated. No changes in brightness or contrast were conducted. A circular area of 20 µm diameter that corresponded to the region of two-photon laser ablation was selected. Following this, a donut-shaped region of width 10 µm was selected surrounding the circular area. This was accomplished using the ‘Edit’ -> ‘Selection’ -> ‘Make Band…’ command in Fiji. This donut-shaped region was added as a region-of-interest (ROI) using ‘Analyze’ -> ‘Tools’ -> ‘ROI Manager…’. The mean gray value for the region was calculated for the time-series prior and post laser-ablation. The intensity values were plotted using R (R Core Team, 2020) version 4.0.3. For comparison, the mean of intensity values of ten frames prior and post laser-ablation were obtained. Comparison was performed using a paired t-test using GraphPad Prism version 9.3.1 for Mac OS X, GraphPad Software, San Diego, California USA, www.graphpad.com.

## Acknowledgements

We thank members of the Singh laboratory for comments on the manuscript, and members of IRIBHM fish facility for technical assistance. We thank M. Martens and J.-M. Vanderwinden from the Light Microscopy Facility for technical assistance at ULB.

## Funding

This work is supported by MISU-PROL funding from the FNRS (40005588) and from Fondation Jaumotte-Demoulin to S.P.S.

## Author Contribution

Conceptualization: S.P.S.; Methodology: I.G.; Resources: M.P.-M., I.G., C.P.; Investigation, Formal analysis, Visualization: M.P.-M.; Writing - Original Draft: M.P.-M., S.P.S.; Supervision, Project administration, Funding acquisition: S.P.S.

## Competing interests

The authors declare no competing or financial interests.

## Data Availability Statement

Raw images and analysis pipeline is available upon request to the corresponding author.

## Supplementary Material

Supplementary Figure 1 along with Legend

Titles and Captions for Movie 1 – 6

## Notes

### Competing Interest Statement

The authors have declared no competing interest.

