## Supplementary Data for "*In vivo* imaging of calcium dynamics in zebrafish hepatocytes"

**Figure S1: Impact of MS-222 (tricaine) on calcium-dynamics in zebrafish hepatocytes** 2

**Movie 1: Impact of Triciane on calcium-dynamics in zebrafish liver** 3

**Movie 2: Representative time-lapse of calcium-dynamics in liver of 5 dpf zebrafish** 3

**Movie 3: Calcium-dynamics in liver of 5, 6 and 7 dpf zebrafish** 3

**Movie 4: Spatially restricted calcium oscillations are induced upon feeding** 3

**Movie 5: Ethanol exposure induces calcium oscillations in zebrafish liver** 3

**Movie 6: Two-photon laser ablation-induced injury induces an increase in cytoplasmic calcium of the neighboring cells** 3

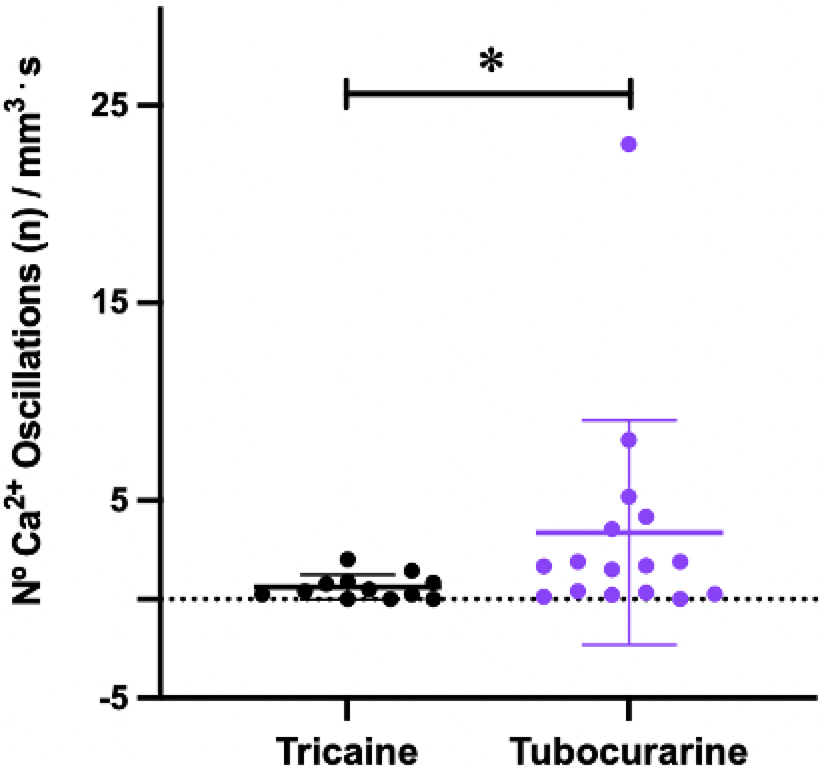


**Figure S1: Impact of MS-222 (tricaine) on calcium-dynamics in zebrafish hepatocytes**

Quantification of calcium oscillations in the liver of 5 dpf *Tg(fabp10a:GCaMP6s)* animals. Animals were either anesthetized in tricaine or immobilized in tubocurarine. Livers of animals treated with triciane had significantly lower calcium oscillations (p-value $< 0.05$, Mann–Whitney U test).

**Movie 1: Impact of Triciane on calcium-dynamics in zebrafish liver**

Representative time-lapse of a liver from *Tg(fabp10a:GCaMP6s)* animals anesthetized in tricaine or immobilized in tubocurarine. Scale-bar: 50 µm.

**Movie 2: Representative time-lapse of calcium-dynamics in liver of 5 dpf zebrafish**

Hepatocytes display repetitive synchronized calcium oscillations. Scale-bar: 50 µm.

**Movie 3: Calcium-dynamics in liver of 5, 6 and 7 dpf zebrafish**

Representative time-lapse images for each stage. Scale-bar: 50 µm.

**Movie 4: Spatially restricted calcium oscillations are induced upon feeding**

*Tg(fabp10a:GCaMP6s)* animals were fed standard rotifer diet from 5 - 7 dpf. At 7 dpf, the time-lapse image was captured. The animal is oriented in the following manner: Dorsal: up; ventral: down; anterior: right; posterior: left. The ventral tip of the liver displays robust calcium oscillations. Scale-bar: 50 µm.

**Movie 5: Ethanol exposure induces calcium oscillations in zebrafish liver**

Representative time-lapse of a liver from 6 dpf *Tg(fabp10a:GCaMP6s)* animals that were either untreated (controls) or treated with 2 % EtOH for 18 - 22 hours. Scale-bar: 50 µm.

**Movie 6: Two-photon laser ablation-induced injury induces an increase in cytoplasmic calcium of the neighboring cells**

A circular region of 20 µm was ablated using two-photon laser ablation. Single-photon calcium imaging was carried out before and after the ablation. A single calcium spike is observed in the neighboring hepatocytes. Scale-bar: 50 µm.
